# Cataloging cysteines in ECOD domains using a protein language model

**DOI:** 10.64898/2026.05.13.724926

**Authors:** Rongqing Yuan, Jesse Durham, Qian Cong, R. Dustin Schaeffer

**Affiliations:** Eugene McDermott Center for Human Growth and Development, University of Texas Southwestern Medical Center, Dallas, TX, USA; Department of Biophysics, University of Texas Southwestern Medical Center, Dallas, TX, USA; Harold C. Simmons Comprehensive Cancer Center, University of Texas Southwestern Medical Center, Dallas, TX, USA

## Abstract

Cysteine is among the most chemically versatile residues in the proteome, existing in three competing functional states: metal coordination, covalent disulfide bonding, and bioactive free thiols. Although these states can be readily assigned from experimentally determined protein structures using simple geometric criteria, accurately annotating them from predicted structures remains challenging. To bridge the gap between predicted structures and functional interpretation, we developed TriCyP (Tri-state Cysteine Predictor), an efficient two-layer neural network built on ESM-2 protein language model embeddings. On an independent benchmark set, TriCyP achieves near-perfect accuracy (AUROC = 0.99) and outperforms existing approaches for predicting both disulfide bonding and metal coordination. We applied TriCyP to classify 2.7 million cysteine residues across 0.9 million ECOD F70 representative domains. The resulting proteome-scale landscape recapitulates established biological patterns. Cysteines are enriched in eukaryotes: disulfide-bonded states are concentrated in extracellular proteins, and metal-coordinating cysteines peak in nuclear proteins owing to the abundance of zinc-finger transcription factors. We further demonstrate the utility of cysteine-state annotation through two pilot studies. First, predicted disulfide-forming cysteines lacking a corresponding structural partner in AlphaFold models may identify either regions of elevated structural uncertainty or latent inter-protein disulfide bonds that stabilize protein-protein interactions. Second, systematic analysis of known and predicted metal-coordinating cysteines across ECOD homologous groups uncovers previously unrecognized metal-binding protein families. This proteome-wide catalog of cysteine states is available as a community resource (http://prodata.swmed.edu/tricyp) and will be integrated into future ECOD releases.

## Introduction

Cysteine is the least abundant of the twenty canonical amino acids, yet it exerts a disproportionate influence on protein structure and function through the distinctive chemistry of its thiol side chain [1]. Its sulfur atom participates in three distinct roles: coordination of metal ions, formation of disulfide bonds, and existence as a free or reactive thiol [2]. These competing fates are determined by the protein and cellular context in which they occur. Because disulfide bonds and metal coordination are often essential for maintaining protein structure and function, these features tend to be highly conserved throughout evolution [3, 4] and can provide valuable information for protein classification, particularly in cysteine-rich small domains and zinc fingers.

The metal-coordinating role of cysteine is ancient and pervasive. Iron-sulfur clusters, among the earliest cofactors in biology, rely on cysteine thiolate ligation for assembly and function in electron transfer, catalysis, and regulatory sensing [5, 6]. Zinc coordination by cysteine is similarly widespread. KRAB C2H2 zinc-finger proteins are the most abundant family of DNA-binding proteins in humans [7]. Beyond structural roles, cysteines can coordinate catalytic metals in a diverse array of enzymes, including alcohol dehydrogenases, sirtuins, metalloproteases, and cytochrome P450s. Comparative analyses of metal-binding proteomes further suggest that zinc-binding sites expanded at the expense of Fe-S binding during eukaryotic evolution, likely driven by rising atmospheric oxygen levels and shifts in trace metal availability in ancient oceans [8–10].

Disulfide bond formation represents a fundamentally distinct functional state of cysteine. In eukaryotic cells, the endoplasmic reticulum (ER) maintains an oxidizing environment in which the ERO1-PDI oxidative folding machinery catalyzes disulfide formation in nascent secretory and membrane proteins [11, 12]. This compartmentalization creates a strong spatial bias: secreted and extracellular proteins are enriched in disulfide bonds, whereas cytoplasmic and nuclear proteins rarely form disulfides under physiological conditions. The structural stability conferred by disulfide bonds has been linked to accelerated sequence evolution in extracellular proteins [13], and enrichment of disulfide-rich secreted proteins in land plants has been associated with adaptation to terrestrial environments [10].

The third cysteine state, the free thiol, has emerged as a regulatory layer of increasingly recognized complexity. The cysteine “redoxome” encompasses thousands of sites subject to reversible oxidative modifications, including sulfenylation, nitrosylation, persulfidation, and glutathionylation [2]. Chemoproteomic profiling across cancer cell lines has revealed extensive heterogeneity in cysteine reactivity, with conserved hyperreactive residues correlating with functional importance [14] and serving as attractive targets for covalent drug discovery [15, 16].

Despite the biological importance of cysteine functional states, proteome-scale annotations of cysteine fate across protein fold space remain unavailable. Previous studies have attempted to annotate disulfide bonds and metal-binding sites in AlphaFold models using geometric criteria [10, 17], but simple Sγ–Sγ proximity within monomeric structures is insufficient for reliable functional assignment of cysteines. For disulfide bonds, AlphaFold frequently positions cysteine pairs within the ∼2.0–2.2 Å distance compatible with covalent bond formation, yet it cannot capture inter-chain disulfides, which constitute approximately 10% of experimentally observed disulfide bonds. Similarly, strong evolutionary covariation within metal-binding motifs, such as zinc fingers, often enables AlphaFold to reproduce the local backbone geometry required to form metal-binding pockets. However, because metal ions are not explicitly incorporated during structure prediction, these pockets frequently collapse, limiting the accuracy of inferred coordination geometries.

Consequently, distinguishing a highly reactive free thiol from a structural disulfide or a latent metal-binding site might require specialized algorithms. Accurate annotation of cysteine states could improve structural modeling—particularly by explicit incorporation of metals in AlphaFold3—and provide important functional insights. Existing metal-binding predictors, however, are restricted to a subset of biologically relevant metals. LMetalSite [18], a sequence-based predictor using ProtT5 embeddings, predicts binding of four free-ion metals (Zn^2+^, Ca^2+^, Mg^2+^, Mn^2+^). Its successor GPSite [19], which adds geometric neural networks operating on predicted structures, retains those four metal channels and adds a heme channel for iron-porphyrin coordination, but like LMetalSite does not address iron-sulfur cluster coordination or other transition metals. The template-based method ZincSight [20] highlights a more fundamental challenge. Although developed to identify zinc-binding sites from coordination geometry, cross-reactivity analyses demonstrated that it detects transition metal-binding sites promiscuously, reflecting the conserved tetrahedral and trigonal coordination geometries shared among Zn, Fe, and Cu sites. Moreover, none of these approaches addresses iron-sulfur cluster coordination, despite iron-sulfur clusters representing the predominant form of cysteine-mediated metal coordination in the PDB.

To comprehensively annotate cysteines involved in both metal coordination and disulfide bond formation in AlphaFold models, we developed TriCyP, a predictor built on ESM-2 protein language model (pLM) embeddings [21]. We reasoned that ESM-2 embeddings encode sufficient evolutionary and biochemical context to infer cysteine functional states directly from sequence, independent of oligomeric state, structure-confidence filters, or metal-type restrictions. Benchmarking and large-scale application of TriCyP to classified domains in the Evolutionary Classification of protein Domains (ECOD) database [22, 23] (**Fig. 1**) demonstrate that this lightweight neural network provides information complementary to structural prediction methods, enabling deeper evolutionary and functional interpretation of AlphaFold models.

**Figure 1.**
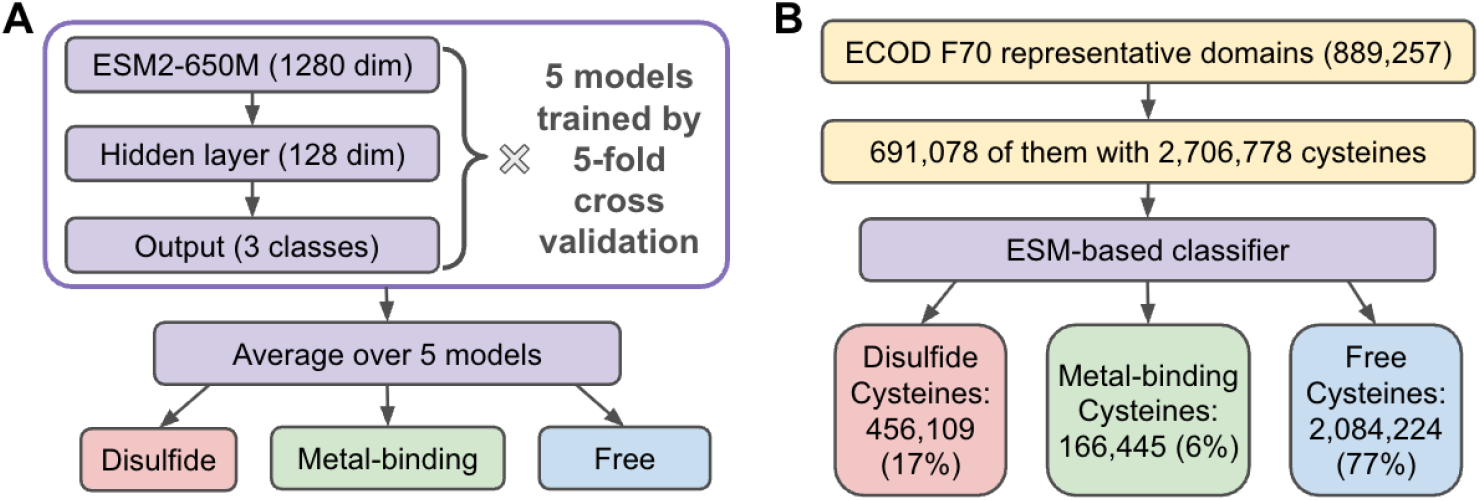
Architecture of TriCyP and its application to ECOD representative domains. **(A)** The TriCyP network takes the 1,280-dimensional ESM-2 embedding vector of a target cysteine residue as input and processes it through a single hidden layer of 128 neurons (164,611 total parameters) to output probabilities for three distinct functional states: disulfide-bonded (Dis), metal-binding (Met), and free thiol (Neg). **(B)** ECOD F70 representative domain sequences were processed through the TriCyP network. Strict probability thresholds (Dis ≥ 0.742, Met ≥ 0.972) were applied to yield high-confidence, discrete functional classifications.

## Results

### Developing a classifier of cysteine states

ESM embeddings have demonstrated significant power in predicting both protein-level and residue-level properties [24–26]. We reasoned that these embeddings could be adapted to predict distinct cysteine functional states by training on experimental structures from the Protein Data Bank (PDB), where cysteine states can be rigorously annotated using geometric criteria. We systematically analyzed 89,698 representative, high-resolution PDB chains (**Fig. S1**), classifying cysteines into three categories: disulfide-forming, metal-binding, and free thiol. Specifically, 100,600 cysteines with an Sγ within 3 Å of another Sγ were classified as disulfide-bonded. Additionally, 8,671 cysteines were identified as metal-binding based on an Sγ distance of less than 2.75 Å from a metal atom (including Zn^2+^, Fe^2+^/^3+^, Ca^2+^, Cu^+^ /^2+^, Hg^2+^, Mg^2+^, Ni^2+^, Cd^2+^, Mn^2+^, and Co^2+^). Within this set, Zn^2+^ and Fe^2+^/^3+^ were the predominant ions, accounting for 66% and 25% of the metal-binding sites, respectively. Only cysteines more than 5 Å from any potential partner were classified as free thiols. A negligible fraction (130) appeared to be both metal-binding and disulfide-forming based on our simplistic criteria and were excluded. Mapping these residues onto the AlphaFold Protein Structure Database (AFDB) entries and performing redundancy reduction resulted in a diverse dataset of 57,353 proteins containing 37,434 disulfide-bonded cysteines, 12,214 metal-binding cysteines, and 151,041 free thiols.

We utilized this extensive dataset to evaluate several network architectures for cysteine classification. These ranged from a streamlined model—converting 1280-dimensional ESM embeddings into state probabilities via a two-layer densely connected neural network (NN)—to more complex geometric neural networks (GNNs) and 3D convolutional neural networks (CNNs) that integrated spatial information from surrounding residues. Surprisingly, we found that these complex, structure-aware architectures did not outperform the simple NN operating solely on the embeddings of the central cysteine. Consequently, we opted for this simple and efficient architecture (**Fig. 1A**). To maximize the utility of our dataset and ensure robust benchmarking, we performed five-fold cross-validation. The final model, TriCyP (Tri-state Cysteine Predictor), is an ensemble of the five models trained during this process.

### Evaluating the performance of cysteine state predictors

We pooled predictions from the test sets of all five TriCyP models to evaluate performance on held-out data across the entire dataset (**Fig. 2A, B, D, E**; red solid lines). We also benchmarked the final ensemble predictor (orange dashed lines), which integrates the five individual models. While the ensemble model exhibits slightly higher performance—likely due to its exposure to the full dataset during various training phases—the held-out performance remains the primary metric for comparison against existing tools. These included the metal-binding predictors GPSite [19] and LMetalSite [18], and the disulfide-detection tool SSBONDPredict [27], which utilizes predicted backbone geometry to score cysteine pairs.

**Figure 2.**
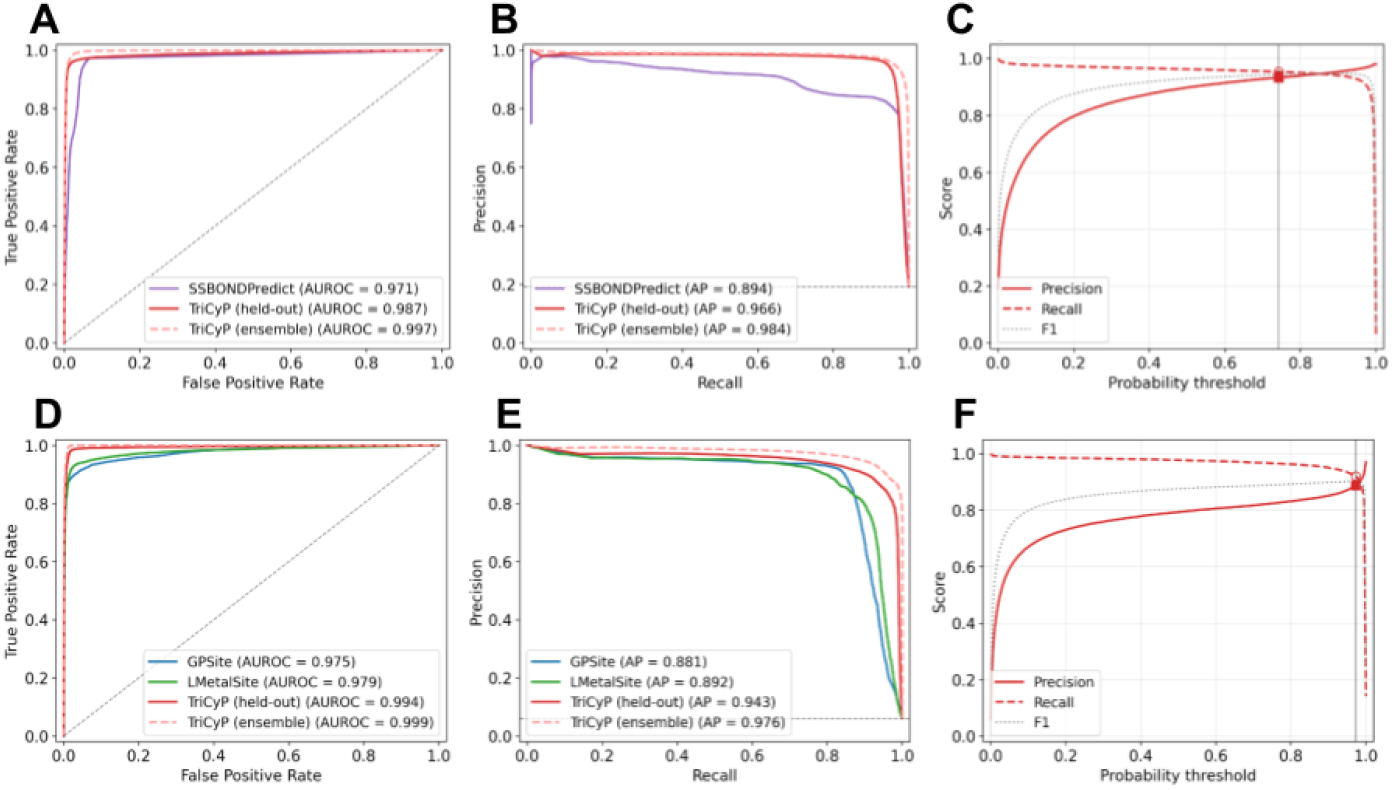
Performance evaluation of cysteine classifiers. Performance for TriCyP is displayed for both held-out cross-validation (solid lines) and the final ensemble model (dashed lines); detailed procedures are provided in the Methods. **(A, D)** ROC curves for **(A)** disulfide and **(D)** metal-binding detection. **(B, E)** Precision-Recall (PR) curves for **(B)** disulfide and **(E)** metal-binding prediction. **(C, F)** Threshold optimization based on TriCyP held-out predictions for **(C)** disulfide and **(F)** metal-binding states. Vertical lines indicate the selected operational thresholds (0.742 for disulfide, 0.972 for metal-binding). Markers (filled squares for precision, open circles for recall) denote performance at these specific thresholds.

For disulfide-bond detection, TriCyP achieved an area under the receiver operating characteristic curve (AUROC) of 0.987 and an average precision (AP) of 0.966 in held-out cross-validation, significantly outperforming SSBONDPredict (AUROC 0.971, AP 0.894) (**Fig. 2A, B**). The diminished performance of SSBONDPredict is primarily driven by a high false-positive rate, likely stemming from the misclassification of metal-binding cysteines as disulfide-bonded. Purely geometric criteria struggle to distinguish a disulfide bridge from a tetrahedral metal-coordination site. In canonical Zn(Cys)_4_ and Fe(Cys)_4_ motifs, such as RING/U-box, rubredoxin, LIM, and zinc fingers, coordinating Sγ atoms are positioned 2.5 – 4 Å apart. This spatial proximity might be misinterpreted by geometric heuristics or iron-biased classifiers as a covalent disulfide bond.

In metal-binding prediction, direct comparisons require nuance; GPSite and LMetalSite are restricted to four metals (Zn^2+^, Ca^2+^, Mg^2+^, and Mn^2+^), whereas TriCyP encompasses all major metal types observed in the PDB. To facilitate comparison, we utilized the maximum predicted probability across all metals supported by GPSite and LMetalSite as their overall metal-binding score. All three tools performed comparably on the shared Zn/Ca/Mg/Mn set, with AUROC values of 0.996, 0.996, and 0.994 for TriCyP, GPSite, and LMetalSite, respectively (**Fig. S2A, B**). However, because GPSite and LMetalSite were not designed to recognize iron-coordinating cysteines—which constitute a non-negligible 14% of metal-binding sites—their performance decays when evaluated against the full diversity of metal sites. Although these tools often assign high probabilities to iron-binding sites due to the similar characteristics between cysteines that coordinate different metals, they ultimately fail to capture the full breadth of iron-sulfur and heme coordination. Consequently, TriCyP (AUROC 0.994) outperformed both GPSite (AUROC 0.975) and LMetalSite (AUROC 0.979) in distinguishing the global metal-binding category (**Fig. 2D, E**) against disulfide bonds or free thiols.

Given its robust multi-class prediction, broader metal coverage, and independence from structural input quality, we selected TriCyP for our proteome-scale cysteine annotations. We determined operational thresholds by maximizing the F1-score (the harmonic mean of precision and recall) for each class (**Fig. 2C, F**), resulting in cutoffs of 0.972 and 0.742, respectively, in recognizing metal-binding and disulfide-bonded cysteines. The stringent threshold for the metal-binding class is necessary due to class imbalance; as metal-binding sites represent only ∼6% of all cysteines, a high cutoff is needed to minimize false positives contributed by other dominating classes.

### Large-scale classification of cysteines in AlphaFold models

We applied TriCyP to F70 (redundant domains showing >70% sequence identity removed) representative domains from the ECOD database, which organizes protein fold space into a hierarchy reflecting evolutionary relationships: Architecture (A-group), Possible Homology (X-group), Homology (H-group), Topology (T-group), and Sequence Family (F-group). ECOD provides a comprehensive catalog of the protein universe, encompassing most PDB entries and AlphaFold models (AFDB) for SwissProt entries and model organisms across diverse phylogenetic lineages. To efficiently survey this space, we analyzed 889,257 representative ECOD domains, applying a 70% sequence identity filter to remove redundancy. Of these, 691,078 domains contained at least one cysteine, totaling 2,706,778 residues (**Fig. 1B**). Applying our optimized probability thresholds (Disulfide ≥ 0.742, Metal ≥ 0.972) classified 456,109 cysteines as disulfide-bonded (17.0%), 166,445 as metal-binding (6.0%), and 2,084,224 as free thiols (77.0%). Prediction confidence was exceptionally high across all functional states, with median probabilities of 0.997 for disulfide-bonded, 0.998 for metal-binding, and 0.961 for free thiol states (**Fig. S3**).

Among these residues, 157,480 originated from PDB entries, allowing for a direct comparison between TriCyP predictions and geometric annotations. Based on geometric criteria, these PDB-derived domains contain 21% disulfide-bonded and 8% metal-binding cysteines (**Fig. 3A**, left). TriCyP predicted a slightly higher fraction of disulfide bonds (24.5%) but a lower fraction of metal-binding sites (5.2%, **Fig. 3A**, middle). The elevated disulfide rate in our predictions likely accounts for cysteines that form inter-domain or inter-protein disulfides not captured by monomeric geometric filters. Conversely, the lower predicted metal-binding fraction reflects a moderate false-negative rate; many true metal-binding sites received high probabilities (around 0.7) but did not pass our stringent 0.972 precision-focused cutoff. Notably, we observed that using isolated domain sequences—rather than full-length protein sequences—to generate ESM embeddings often deflated the predicted probabilities. This suggests that ESM-2 embeddings do not merely reflect the local properties of a single residue, but are deeply influenced by the broader sequence context and surrounding environment. For optimal performance, we recommend that TriCyP users utilize full-length protein sequences.

**Figure 3.**
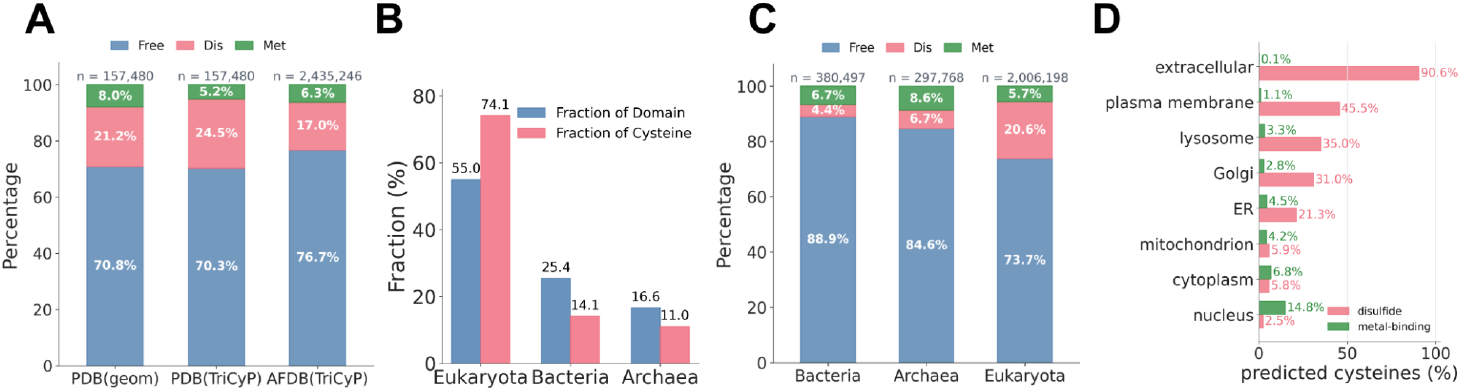
Distribution of metal-binding and disulfide-bonded cysteines across structural sources, domains of life, and subcellular compartments. **(A)** Comparative analysis of cysteine functional states by structural source. Cysteines in PDB-sourced domains are annotated using both geometric criteria (geom) and TriCyP predictions; cysteines in AFDB-sourced domains are annotated by TriCyP alone. Stacked bars represent the fraction of cysteines classified as free thiol (Free, blue), disulfide-bonded (Dis, red), or metal-binding (Met, green). **(B)** Relative abundance of ECOD representative domains (blue) compared to total cysteine counts (red) for Eukaryota, Bacteria, and Archaea, highlighting the disproportionate cysteine enrichment in eukaryotes. **(C)** TriCyP-based functional classification across the three domains of life. Stacked bars illustrate free thiol (blue), disulfide (red), and metal-binding (green) fractions. While metal-binding rates remain comparable across domains, disulfide rates show significant divergence. **(D)** Eukaryotic cysteine classification by subcellular localization. Disulfide rates (red) recapitulate the oxidative folding gradient, declining from secreted proteins (91%) through the ER (21%) to the cytoplasm (5.8%) and nucleus (2.5%). Conversely, metal-binding rates (green) peak in the nucleus (14.8%) and cytoplasm (6.8%)—where zinc-finger transcription factors and RING/U-box E3 ligases are concentrated—and decline throughout the secretory pathway (0.1% in secreted proteins).

The much larger set of AFDB-sourced domains exhibited a lower disulfide rate (17.0%) compared to the PDB set (21.0%, **Fig. 3A**, right), likely reflecting the broader, less-biased fold coverage of AFDB models. Proteins containing disulfide bonds are often more rigid and stable, making them easier targets for experimental structure determination—a bias that likely explains the higher disulfide density in the PDB-deposited set.

Across the three domains of life, Eukaryotes account for 55% of all representative domains but contribute a disproportionate 74% of all cysteines. In contrast, despite constituting a significant portion of the protein universe, Bacteria and Archaea contribute only 14% and 11% of total cysteines, respectively (**Fig. 3B**). Eukaryotic domains are thus disproportionately cysteine-rich. Disulfide bonding is also markedly higher in Eukaryotes (20.6%) compared to Bacteria (4.4%) and Archaea (6.7%) (**Fig. 3C**). This disparity is driven by the fact that many eukaryotic proteins fold within the ER, a specialized, oxidizing environment designed for bond formation [28, 29] —in contrast to the highly reducing cytoplasm where the majority of prokaryotic folding occurs [30]. This structural “stapling” via disulfides is essential for stabilizing the complex secreted and membrane-associated proteins that multicellular organisms require to navigate harsh extracellular environments [4].

The sophisticated compartmentalization of eukaryotic cells establishes distinct chemical environments that allow for a sharp subcellular oxidative folding gradient [31]. This biological transition—moving from the reducing interior of the nucleus and cytoplasm to the oxidizing conditions of the secretory pathway [32]—is remarkably recapitulated by TriCyP from sequence alone. In reducing compartments, high concentrations of antioxidants like glutathione and thioredoxin prevent cysteine oxidation [33], whereas the ER and extracellular space provide the specialized machinery for covalent bond formation.

TriCyP-predicted disulfide rates mirror this biological architecture: rates decline from 91% in extracellular proteins through the plasma membrane (46%), lysosome (35%), Golgi (31%), and ER (21%), reaching a minimum in the mitochondrion (5.9%), cytoplasm (5.8%), and nucleus (2.5%) (**Fig. 3D**). Metal-binding patterns exhibit the inverse trend, peaking in the nucleus (14.8%) and cytoplasm (6.8%)—compartments where zinc-finger transcription factors and RING/U-box E3 ligases are concentrated—and declining through the ER (4.5%) to extracellular proteins (0.2%). These results demonstrate that TriCyP successfully captures both the oxidative folding gradient and the subcellular distribution of metal-coordinating motifs using sequence information alone.

### TriCyP predictions reveal cysteines participating in inter-protein interactions

The superior performance of TriCyP over geometry-based methods like SSBONDPredict suggests that sequence-based predictors can be more reliable for disulfide detection than relying solely on AlphaFold models. We reasoned that evolutionary information captured by ESM-2 embeddings could be integrated with AlphaFold structures to provide deeper structural insights. To test this, we analyzed the minimal Sγ-Sγ distance in PDB structures and AlphaFold models for all predicted disulfide-bonded cysteines. While the majority exhibit distances below 3 Å—clustering near the canonical disulfide bond length of 2.03 Å—approximately 10% lack a nearby cysteine partner within the domain **(Fig. 4A**). While some of these cases may represent false positives, others likely highlight cysteines that form disulfide bonds with partners not included in the single domains.

**Figure 4.**
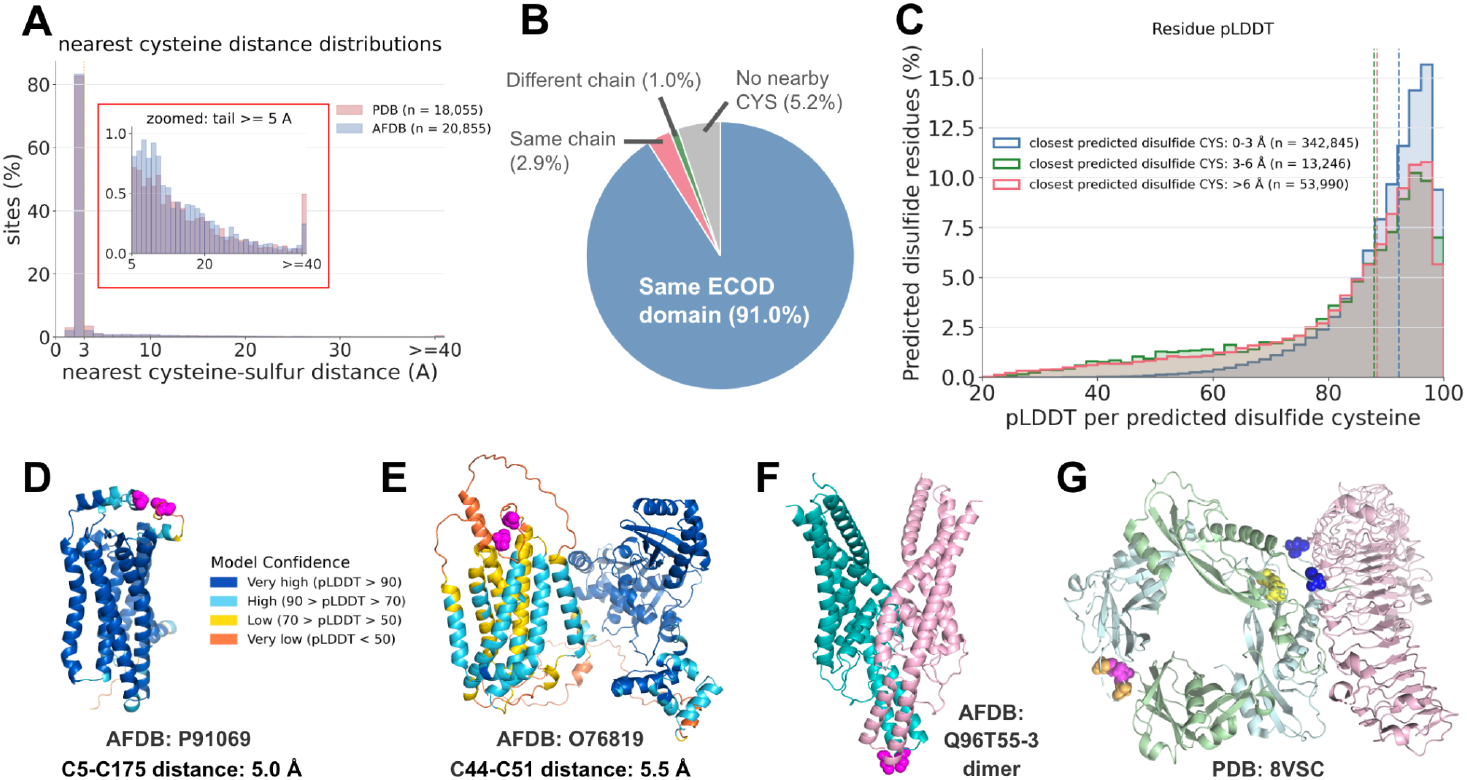
Structural validation of TriCyP predictions and identification of inter-protein disulfides. **(A)** Distribution of minimal Sγ–Sγ distances for cysteines predicted as disulfide-bonded in PDB experimental structures (red, n = 18,055) and AlphaFold models (blue, n = 20,855). Both distributions exhibit a sharp peak at ∼2.03 Å, the canonical disulfide bond length. **(B)** Classification of predicted disulfide-bonded cysteines in PDB structures based on their biological context: intra-domain, inter-domain, or inter-chain. **(C)** Predicted disulfide cysteines with a nearby predicted partner exhibit significantly higher average pLDDT scores than those lacking a proximal partner. **(D-E)** Representative examples of predicted disulfide cysteines (magenta spheres) characterized by distorted geometries and low pLDDT scores in AlphaFold models. These regions highlight structural mismodeling where AlphaFold likely failed to resolve a covalent bond; structures are colored by pLDDT to indicate local model quality. **(F-G)** Instances where a predicted disulfide-bonded cysteine lacks an intra-chain partner in the monomer but forms an inter-chain bridge in the oligomeric complex. **(F)** KCNK16 dimer illustrating the predicted inter-chain disulfide bridge (magenta spheres). **(G)** TGFB1 dimer (pale cyan and pale green) in complex with LRRC32 (pale pink). Inter-protein disulfide cysteines successfully identified (probability > 0.742) by TriCyP are shown as magenta and blue spheres. Additional inter-protein disulfides that received moderate or low probabilities are indicated by orange and yellow spheres, respectively.

Analysis of PDB-derived domains within their full biological assemblies supports this hypothesis: 2.9% of predicted disulfide-bonded cysteines are covalently linked to cysteines in other domains, while 1.0% are bonded to cysteines from separate protein chains (**Fig. 4B**). A recent PDB-wide survey reported that approximately 5% of all disulfide bonds in deposited structures bridge two protein chains[34], a frequency significantly higher than that currently identified by TriCyP. This disparity suggests that while TriCyP is capable of detecting inter-protein signals, it inherently prioritizes the intra-protein context. More robust detection of these inter-chain covalent linkages will likely require the joint analysis of multiple protein sequences or the implementation of complex-aware architectures that can explicitly model the interface between subunits.

Compared to PDB-derived domains, AFDB-derived domains contain a higher fraction of predicted disulfide cysteines without a nearby partner (see the distribution tail inside the red box in **Fig. 4A**). We hypothesized that this discrepancy results from disulfide bonds being poorly modeled during AlphaFold’s structural prediction. In support of this, we found that predicted disulfide cysteines lacking a partner systematically exhibit lower pLDDT scores (**Fig. 4C**). **Figures 4D** and **4E** illustrate potential examples where TriCyP correctly identifies a disulfide state for two cysteines that AlphaFold positioned in proximity but failed to close into a covalent bond.

Despite current limitations, TriCyP’s ability to recognize inter-protein disulfide bonds inspires its application in assisting oligomer modeling. Strong predictions without intra-protein partners can serve as markers for inter-protein interfaces. We manually examined several proteins known to form inter-protein disulfide bonds, such as the potassium channel subfamily K member 16 (KCNK16). KCNK16 forms a covalent dimer through a disulfide bridge at the subunit interface [35, 36]. TriCyP correctly predicted the disulfide state for residue 60 (**Fig. 4F**, magenta spheres), which indeed forms an inter-protein bond in the AlphaFold model of the oligomeric structure.

Transforming growth factor beta-1 (TGFB1) provides another example; the experimental structure of TGFB1 with its activator LRRC32 reveals three inter-protein disulfide bonds (**Fig. 4G**, magenta, orange, and yellow spheres) between the TGFB1 monomers and additional bonds linking them to LRRC32 (**Fig. 4G**, blue spheres). However, not all these bonds were strongly predicted by TriCyP—some were classified as free thiols or received moderate probabilities below our stringent cutoff. Because ESM-2 was trained primarily on monomeric sequences and our embeddings are generated from monomers, the signal for inter-protein disulfide bonds may be attenuated. Nevertheless, the presence of an “orphaned” high-confidence disulfide prediction remains a powerful indicator of potential oligomerization or structural mismodeling.

### Distribution of metal-binding cysteines across ECOD H-groups

We analyzed the distribution of metal-binding cysteines across 2,194 ECOD H-groups that met two criteria: (1) containing at least 10 F70 representative domains and (2) including both PDB-sourced and AFDB-sourced representatives. For PDB-sourced domains, metal-binding status was determined by geometric criteria, while AFDB-sourced domains were annotated using TriCyP. Of the 889,257 representative ECOD domains, 53,460 (6.0%) were predicted to bind metals; however, these are distributed highly unevenly across the ECOD hierarchy. Only 542 H-groups contain metal-binding cysteines.

**Figure 5A** lists the H-groups containing the largest absolute number of metal-binding domains. Because these groups were selected by count rather than percentage (enrichment), they fall into two distinct biological categories. Roughly half of these H-groups (highlighted in green, **Fig. 5A**) consist of dedicated metalloregulatory or redox-active families, such as Zinc Fingers and 4Fe-4S Ferredoxins. In these groups, metal coordination is a near-universal feature, and the large count reflects both the group’s size and its high internal percentage of metal-binding members. The remaining half (highlighted in red, **Fig. 5A**) represents a different phenomenon. These are expansive, ubiquitous superfamilies—such as P-loop domains, Rossmann folds, and TIM barrels—where metal-binding cysteines are present in only a small fraction of the total membership. However, because these H-groups contain thousands of members, even a low-frequency feature translates into a large absolute number of metal-binding domains. In many of these cases, the metal-binding motif acts as a specialized regulatory or catalytic site. Interestingly, many of these motifs appear to be autonomous functional units that the automated ECOD pipeline failed to partition as separate domains. Explicitly identifying these “buried” metal-binding sites through TriCyP could significantly improve the precision of future domain parsing and hierarchical assignments.

**Figure 5.**
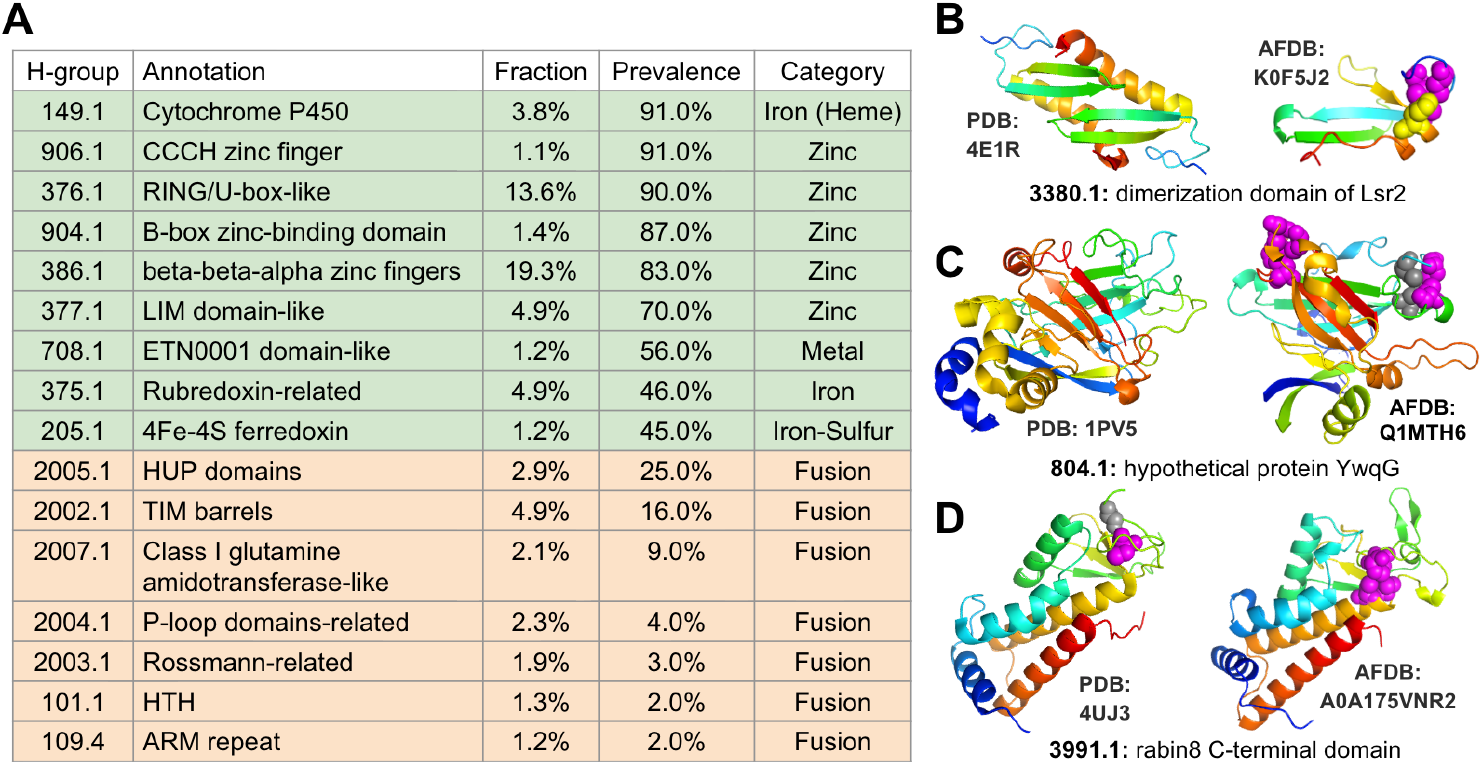
Analysis of metal-binding domains across ECOD H-groups. **(A)** H-groups containing the highest absolute counts of predicted metal-binding domains. Groups highlighted in green represent dedicated metalloregulatory or redox-active domains; groups in red represent large, ubiquitous superfamilies where metal-binding motifs occur as specialized functional or regulatory elements. **(B–D)** Three previously uncharacterized metal-coordinating H-groups identified by TriCyP, each comparing a PDB-sourced representative (left) with a corresponding AFDB-sourced representative (right). H-group IDs and H-group names are provided below each panel, with specific protein accessions labeled by the 3D structures. Magenta spheres indicate high-confidence TriCyP-predicted metal-binding cysteines. Grey spheres indicate sub-threshold cysteines (probability > 0.7) proximal to the predicted site that likely contribute to the coordination environment. Yellow spheres indicate histidines positioned to serve as co-ligands within the metal-binding site.

Beyond refining established groups, we utilized TriCyP to identify H-groups that likely function as dedicated metal coordinators but have previously escaped structural characterization. We identified three specific H-groups where more than half of the AFDB-sourced representatives contain predicted metal-binding cysteines, yet the corresponding PDB entries lacked any geometric evidence of metal binding (**Fig. 5B–D**). In the Lsr2 dimerization domain (H-group 3380.1), YwqG hypothetical protein (H-group 804.1), and Rabin8 C-terminal domain (H-group 3991.1), TriCyP identified high-confidence metal-binding cysteines (magenta spheres). These “anchor” residues are frequently flanked by other high-probability coordinating cysteines (grey spheres) or histidines (yellow spheres) that, while falling just below our stringent classification threshold, likely serve as essential co-ligands to complete the metal-binding site.

Detailed inspection suggests that the available experimental structures for these groups represent instances where the metal-binding site was absent in the specific family members that have been experimentally studied. For example, while the experimental Rabin8 structure retains two potential coordinating cysteines, it lacks the full four-cysteine motif observed in its AFDB-sourced homologs. These cases illustrate how structural databases can miss entire classes of metal-binding folds when the only available experimental members are degenerate paralogs or artifacts of a traditionally metal-active family.

## Discussion

### Advantage and limitations of protein language models

It is remarkable that TriCyP—a relatively simple model built upon the ESM-2 embedding of a single position—can achieve near-ideal performance in predicting disulfide bonds and metal-binding sites. During development, we explored more sophisticated architectures designed to explicitly incorporate broader residue context, yet these models yielded no performance gains over our simpler approach. This observation is further supported by the comparable performance of existing tools such as GPSite and LMetalSite. While LMetalSite, like TriCyP, leverages pLM embeddings, GPSite [19] employs a complex, edge-enhanced graph neural network that integrates backbone and side-chain coordinates, DSSP features, and inter-residue distances within a 15 Å radius. The convergence in performance between these tools suggests that the inclusion of structural features from AlphaFold models does not improve metal-binding prediction. It appears that the essential information required to identify these functional states is already captured within the embedding of a single position.

Furthermore, because AlphaFold models often lack the necessary precision in ligand-binding pockets or disulfide geometries, incorporating 3D structural data from these models can increase computational complexity without a concomitant improvement in accuracy. Willems et al. [10] reported that AlphaFold occasionally forms spurious disulfide bonds between cysteines that coordinate iron-sulfur clusters when they are positioned in close proximity in the absence of a cluster. This is a clear instance of the coordination geometry degeneracy documented by Mechtinger et al. in ZincSight, where tools trained on zinc-binding sites exhibit cross-reactivity with other metal types [20]. Such structural artifacts likely explain why GPSite’s own ablation studies show a performance degradation when using predicted structures compared to experimental structures (average AUPR 0.594 vs. 0.656) [19].

The superb performance of pLM embeddings in this context stems from the fact that disulfide and metal-binding sites are highly conserved and vital for structural and functional integrity. Consequently, pLMs learn to recognize these patterns as they are highly informative for recovering masked tokens during the self-supervised training process. However, caution is required when utilizing pLM-based predictors. For instance, we observed a decrease in performance when utilizing isolated domain sequences rather than full-length protein sequences to generate embeddings. Because pLM embeddings for a single position integrate contextual information from the entire input sequence, and TriCyP was trained on full-length data, using truncated domain sequences can compromise prediction accuracy.

Additionally, pLM-based models are inherently limited in their ability to predict functional states gained through mutations. Because pLMs represent a “statistical average” of evolutionary success rather than a physical engine for atomic interactions, they do not explicitly calculate bond energies or the precise spatial distances between sulfur atoms. Consequently, if a new cysteine is introduced via mutation, the model may treat it as a generic residue or an “evolutionary error” rather than recognizing the formation of a novel covalent bridge or coordination site.

### Comparison with prior proteome-scale cysteine annotation

The most directly comparable prior work is that of Willems et al. [10], who utilized AlphaFold2 predicted structures to annotate cysteine disulfides and metal-binding sites across fifteen plant proteomes. Their approach was entirely structure-based: disulfides were identified by Sγ-Sγ distances < 2.5 Å, and metal sites were detected using a geometric template-matching algorithm that superimposes known coordination geometries onto predicted structures. Their findings parallel ours in several respects, particularly the observation that disulfide rates are strongly compartmentalized. They reported that 58% of cysteines in secretory pathway proteins form disulfides compared to 3% in mitochondrial proteins—a trend consistent with the gradient we observed across all eukaryotes (91% in secreted proteins vs. 5.9% in mitochondrial proteins). The quantitative differences between our results likely reflect the broader scope of our study, which spans all domains of life rather than being restricted to plants.

Our work extends the analysis of Willems et al. in three significant ways. First, we have developed, benchmarked, and released TriCyP as a standalone predictor, enabling the community to apply this analysis to any protein sequence. Second, by assigning functional labels to individual cysteines rather than relying on Sγ-Sγ proximity, we circumvent the “metal-disulfide conflation” documented by Willems et al. In their study, they identified 960 instances where AlphaFold positioned iron-sulfur-coordinating cysteines at disulfide-like distances, causing their geometric pipeline to misidentify them as covalent bonds. The three-state TriCyP classifier correctly labels these residues as metal-binding directly from sequence context. Furthermore, because TriCyP is agnostic to the oligomeric state, it can detect inter-chain disulfides that remain invisible to structural predictors focused on monomers. Third, by analyzing the full ECOD representative set rather than individual proteomes, we can examine cysteine function across the entirety of known protein fold space. This approach reveals evolutionary patterns that are invisible within a single kingdom, such as the divergence in disulfide usage between Bacteria and Eukaryota within homologous groups, as well as the conservation of metal-binding sites across diverse folds.

### Limitations of our cysteine state predictor

Despite the impressive performance of TriCyP in annotating cysteines, there are many key limitations that need to be addressed in the future. First, the three-state classification treats cysteine fate as a static, intrinsic property of the protein sequence. In reality, the cysteine functional state is context-dependent and dynamic. Allosteric disulfides form and break as part of protein regulation [37], zinc-binding cysteines can release metal and form disulfides under oxidative conditions [38], and redox-active cysteines in thioredoxin family enzymes cycle between thiol and disulfide states as part of their catalytic mechanism. The classifier cannot account for cofactor availability: a protein whose iron-sulfur cluster has not yet been assembled by the ISC/SUF machinery [39] has free cysteines at positions the classifier labels as metal-binding. TriCyP is trained to assign each cysteine to the dominant state represented in the training data consisting of PDB entries, which captures a single snapshot of a potentially dynamic system.

Second, TriCyP classifies cysteines into three broad categories, but cysteines within a category could play quite different roles. The metal-binding class does not distinguish between metal types or between direct ion coordination and cofactor-mediated coordination. The free thiol class is particularly heterogeneous. It includes structurally buried inert cysteines, solvent-exposed reactive thiols subject to oxidative post-translational modifications, and catalytic nucleophiles in enzymes such as cysteine proteases and phosphatases. These are functionally distinct categories unified only by the absence of a disulfide bond or metal coordination. Distinguishing reactive free cysteines from non-reactive cysteines is the subject of ongoing work in our lab.

Third, although TriCyP has the ability to predict inter-protein disulfide bonds, its recall in such disulfide bonds is much lower. Among disulfide bonds predicted by TriCyP, only about 1% are inter-protein based on our analysis of PDB entries, missing the majority of such inter-protein disulfide bonds, which is expected to constitute 5% of disulfide bonds in PDB. However, inter-protein disulfide bonds might be critical for stabilizing both homo-oligomers and hetero-oligomers, and effectively predicting them would require coupling between disulfide bond prediction and protein complex modeling.

### Incorporating residue feature predictions into future ECOD development

As the developers of ECOD, we propose that annotating specialized cysteines provides a granular, compositional view of biochemical function that enhances traditional domain-level function assignments. Rather than attributing a single, broad function to an entire domain, cysteine classification captures specific residue-level chemistry, illustrating how these functional “anchors” are distributed within and across evolutionary families. Instead of trying to fit complex proteins into rigid, pre-defined categories, we look at the specific jobs of individual amino acids. This allows us to map out a protein’s function piece-by-piece based on its actual chemistry.

Furthermore, the annotation of disulfide bonds and metal-binding cysteines is instrumental for recognizing remote homology and resolving persistent ambiguities in domain classification. While our automated pipelines effectively partition AlphaFold models into globular domains [40] and assign them within the ECOD hierarchy [41], small cysteine-rich domains frequently evade these automated workflows [42]. By predicting disulfide connectivity and metal-coordination sites, we can identify conserved structural frameworks and shared biochemical signatures that persist even when sequence identity and structure similarity have diverged beyond the detection limits of traditional alignment methods.

## Methods

### Training and testing dataset preparation

We retrieved all protein sequences from PDB entries released prior to June 2025 and clustered them using MMseqs2 [43] (95% identity and 95% coverage thresholds). From each cluster, the chain with the highest resolution was selected as the representative. We excluded entries lacking resolution data (e.g., those determined by NMR) or those with a resolution poorer than 5 Å. Using the structural coordinates of these representative chains, we classified cysteines into three functional categories based on their local chemical environment:

1. **Disulfide-bonded (Dis):** Defined by an Sγ to Sγ distance of less than 3 Å.
2. **Metal-binding (Met):** Defined by an Sγ to metal ion distance of less than 2.75 Å for a specific set of ions (Zn^2+^, Fe^2+^/^3+^, Ca^2+^, Cu^+^/^2+^, Hg^2+^, Mg^2+^, Ni^2+^, Cd^2+^, Mn^2+^, and Co^2+^).
3. **Free thiols (Neg):** Defined by an Sγ distance of greater than 5 Å from any other cysteine or metal ion.

To mitigate the impact of structural imperfections, we implemented a “grey zone” for cysteines located within 5 Å of a potential partner that failed to meet the strict bonding criteria; these residues were excluded from the training set. Additionally, we excluded the rare instances where a cysteine met the criteria for both disulfide and metal binding to ensure distinct label assignment.

We kept PDB chains that contain cysteines in any of these three categories and we mapped them to AFDB (version4) entries by MMseqs, requiring the hit to show sequence identity over 95% to the query and coverage over 90% of the query residues. For each mapped cysteine, we further checked residues that are less than 10Å away from it and only kept cysteines whose neighboring residues are identical between the query PDB sequence and the hit AFDB entry. We transferred the cysteine category labels in PDB chains to the AFDB residues they map to. We performed this mapping because we were planning to use the predicted 3D structures in our predictor. Because one PDB chain might map to multiple AFDB entries, and thus we again removed redundancy in the AFDB entries through clustering by MMseqs2 (90% identity and 25% coverage threshold), and the representative protein from each cluster were taken to include in our training/testing dataset.

We retained PDB chains containing cysteines from all three functional categories and mapped them to AFDB entries using MMseqs2. To ensure high-confidence matches, we required a sequence identity >95% and a coverage threshold of >90% relative to the query PDB sequence. For each mapped cysteine, we performed a secondary validation by inspecting all neighbor residues within a 10 Å radius; only cysteines whose local neighborhoods were identical between the PDB and AFDB entries were kept. This stringent mapping was performed to ensure that the mapped cysteine in AFDB models will fall into the same functional category. To address cases where a single PDB chain mapped to multiple AFDB entries and contributed unproportionally to the final dataset, we applied a final redundancy reduction step using MMseqs2 clustering (90% identity and 25% coverage thresholds). The representative protein from each resulting cluster was selected for inclusion in our final training and testing datasets.

### Building and training the TriCyP classifier

Each protein sequence in the training/testing dataset was processed through the pretrained ESM-2 650M language model (esm2_t33_650M_UR50D) [21] to generate per-residue embeddings. The 1,280-dimensional vector from the final transformer layer at each cysteine position served as the input features. The TriCyP classifier is a two-layer neural network consisting of an initial linear layer (1,280 to 128 units), batch normalization, ReLU activation, and dropout, followed by a final linear layer (128 to 3 classes), totaling 164,611 trainable parameters. We implemented a five-fold cross-validation strategy, randomly partitioning the proteins into five equal folds. For each iteration, a model was trained on four folds (∼80% of proteins) while the fifth was held out for evaluation. During training, a subset of the training data (25%) was reserved as a validation set to monitor progress and perform early stopping based on optimal validation loss.

Models were trained using the Adam optimizer with an L2 weight decay of 1×10^-5^ for regularization. We conducted an exhaustive hyperparameter grid search covering hidden layer sizes (64, 128, 256), learning rates (0.00001, 0.00002, 0.00004), batch sizes (64, 128, 256), and dropout rates (0.2, 0.4, 0.6). To address the inherent class imbalance among cysteine states, we evaluated three strategies: (1) weighted cross-entropy loss with class weights inversely proportional to frequency, (2) focal loss (gamma=2.0) combined with class-specific weights, and (3) balanced sampling via a weighted random sampler to ensure equitable representation of minority classes during each epoch.

We conducted an exhaustive grid search across all hyperparameter combinations, evaluating model performance on the validation set using a one-vs-rest AUROC approach. Possibly due to that this is a straightforward task, the choice of hyperparameters do not affect the validation performance much, and we thus did not perform further hyperparameter search and just used the following best-performing set of hyperparameters: number of hidden layers: 128, learning rate: 0.00004, batch size: 64, dropout rate: 0.2, use class-specific weights for cross entropy loss but do not use focal loss, use balanced sampling of different classes of cysteines.

The five models trained through this cross-validation process were integrated into an ensemble, where the final prediction for each cysteine represents the average of the five predicted probabilities. At the inference stage, rather than defaulting to the *argmax* class, we applied optimized probability thresholds to balance classification precision and recall: a cysteine is classified as a disulfide if P(disulfide) ≥ 0.742 and as metal-binding if P(metal-binding) ≥ 0.972; otherwise, it is classified as a free thiol. These specific thresholds were selected to maximize the F1-score for disulfide and metal-binding cysteine predictions, respectively, determined by analyzing the pooled predictions from the held-out sets of all five models.

### Evaluating the performance of cysteine classifiers

To assess model generalization, we compared ensemble predictions (where all five models scored the entire dataset) against out-of-fold predictions (where each protein was scored strictly by the model that excluded it during training). The out-of-fold metal-binding AUROC (0.994) was marginally lower than the ensemble AUROC (0.999), indicating only modest overfitting. To ensure a rigorous assessment, all subsequent benchmark evaluations utilized the out-of-fold predictions. Overall performance for both metal-binding and disulfide detection was quantified using the AUROC and AP, evaluated against all other cysteine categories.

We benchmarked TriCyP against three specialist predictors using the comprehensive dataset derived from PDB structures mapped to AFDB version 4 sequences. Because some tools require 3D coordinates and AFDB v4 structures are no longer publicly hosted, we utilized AFDB v6 structures for this evaluation. To account for versioning discrepancies, we excluded 5,721 proteins (10.3%) that were obsolete in the v6 release and 1,001 proteins (1.8%) exhibiting sequence mismatches between v4 and v6. This filtering yielded a shared evaluation set of 173,241 cysteines across 48,659 proteins, comprising 129,009 free thiols, 33,541 disulfides, and 10,691 metal-binding cysteines.

The baseline tools for metal-binding prediction included LMetalSite, which features four independent binary classifiers for zinc (Zn), calcium (Ca), magnesium (Mg), and manganese (Mn); and GPSite, a multi-task tool that generates predictions for the same four metals in addition to heme. To establish a standardized primary metric for these two tools, we assigned each cysteine the maximum predicted probability across the four mutually supported metal types (Zn, Ca, Mg, Mn). GPSite’s performance on its specific heme (iron-porphyrin) channel is reported separately in supplementary **Figure S2**. Because GPSite and LMetalSite are restricted to these specific ions and cannot capture the full spectrum of metalloproteins, we stratified the benchmark evaluations by metal type to ensure an equitable comparison. To achieve this, every metal-binding cysteine in the dataset was mapped back to its source PDB structure to explicitly identify the coordinating metal ion.

Finally, for disulfide benchmarking, we utilized SSBONDPredict, a structure-based tool that evaluates cysteine pairs based on backbone and Cβ geometry. To adapt its pairwise output for our per-residue evaluation, each individual cysteine was assigned the maximum score derived from any predicted disulfide pair in which it was involved.

### Annotating cysteines in representative ECOD domains and subsequent analysis

We utilized the ECOD database’s 70% sequence identity clustering (F70) to select representative domains, yielding a total of 889,257 representatives. Of these, 691,078 domains contained at least one cysteine residue, providing a dataset of 2,706,778 total cysteines. We extracted the corresponding domain sequences, computed per-residue ESM-2 embeddings, and applied TriCyP to classify each cysteine. Where available, experimental structural evidence—derived from PDB LINK/STRUCT_CONN records for metal coordination and an intra-domain Sγ-Sγ distance threshold of < 2.5 Å for disulfides—was recorded as corroborating annotation. However, this empirical data was used purely for reference and did not override the TriCyP predictions. Importantly, because these structural annotations are strictly domain-scoped, they inherently exclude inter-domain, inter-chain, and unassigned-partner disulfide bonds.

UniProt accessions were mapped for 653,528 classified domains (representing 94.6% of the cysteine-containing F70 representatives). Subcellular location annotations were retrieved via the UniProt REST API for 107,191 of these proteins. The free-text location fields were then systematically parsed into simplified subcellular compartment categories (Extracellular, Plasma membrane, ER, Golgi, Lysosome, Mitochondrion, Cytoplasm, and Nucleus). Taxonomy assignments for each protein were sourced from the ECOD database, which categorizes entries by superkingdom (Eukaryota, Bacteria, Archaea, Viruses) based on NCBI taxonomy.

### Use of large-language models

Large-language models (e.g., ChatGPT 5.3, Claude Opus 4.7, Claude Sonnet 4.6) were used in the derivation and reporting of this work. The ESM-2 three-state cysteine classifier, model architecture, and threshold calibration were not guided by LLMs. LLM assistance was used to aid in writing analysis and visualization scripts, generating documentation, and revising manuscripts for clarity and concision. All scientific ideas, hypotheses, experimental design, results interpretation, and conclusions presented in this work originate with the authors. The authors take full responsibility for the content of the manuscript and have verified all reported analyses and citations.

## Software and data availability

A browser for the per-domain cysteine annotations is available at prodata.swmed.edu/tricyp. A Github repository for the TriCyP predictor is available at https://github.com/conglab2020/tricyp.Code, trained model weights, per-cysteine predictions for all ECOD F70 representative domains, the benchmark dataset with predictions from various tools, per-domain structure shards, and an SQL dump of the analysis schema are deposited on Zenodo at doi.org/10.5281/zenodo.20072069.

