## Supplementary figures and images for "Cataloging cysteines in ECOD domains using a protein language model"

### Figure S1

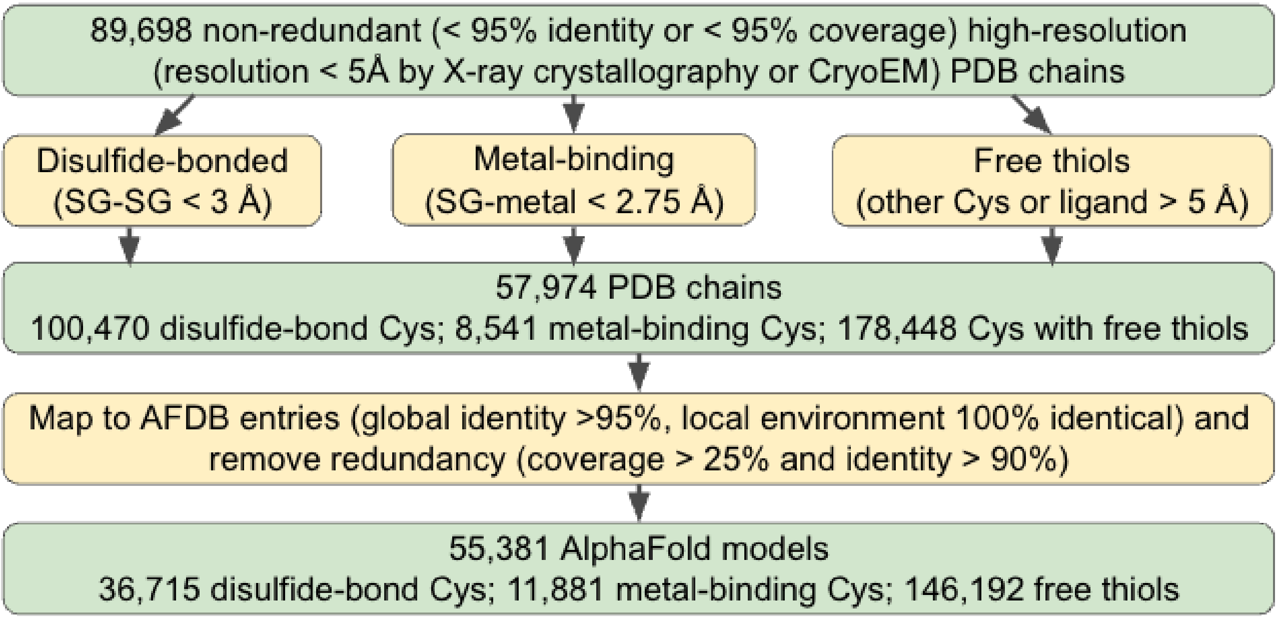


**Figure S1. The pipeline to prepare training and benchmark datasets for TriCyP.**
