## Supplementary material for "Cataloging cysteines in ECOD domains using a protein language model": Figure S2

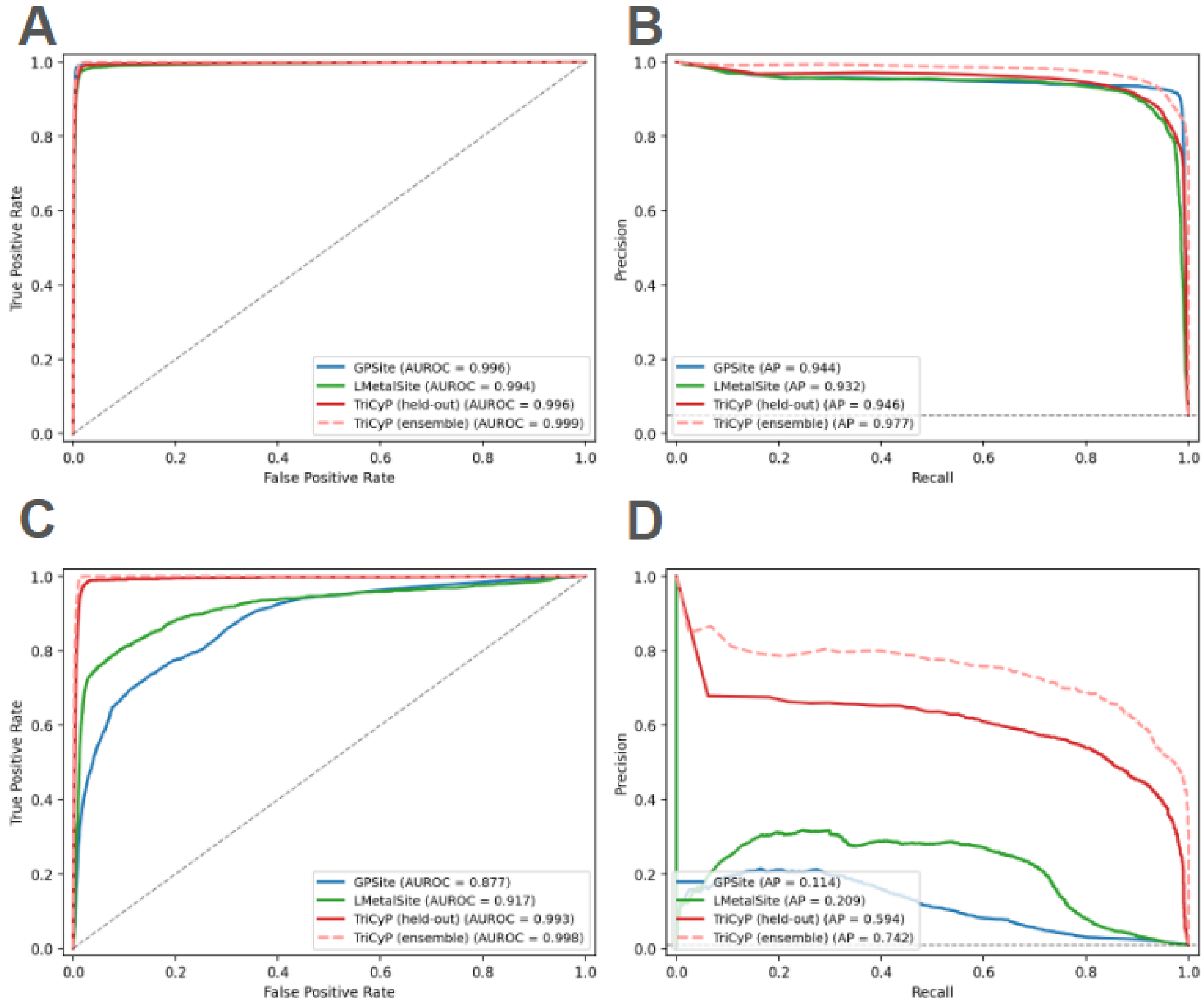


**Figure S2. Performance comparison of TriCyP against existing predictors across diverse metal-binding states. (A–B)** Comparative performance metrics for zinc (Zn), calcium (Ca), magnesium (Mg), and manganese (Mn)—the specific metal types currently supported by existing state-of-the-art tools. **(C–D)** Evaluation of iron (Fe) binding prediction. Iron is a ubiquitous biological cofactor that is frequently coordinated by cysteines (e.g., in Fe-S clusters and heme groups), yet it remains largely unsupported by previous sequence-based prediction frameworks.
