## Supplementary material for "Cataloging cysteines in ECOD domains using a protein language model": Figure S3

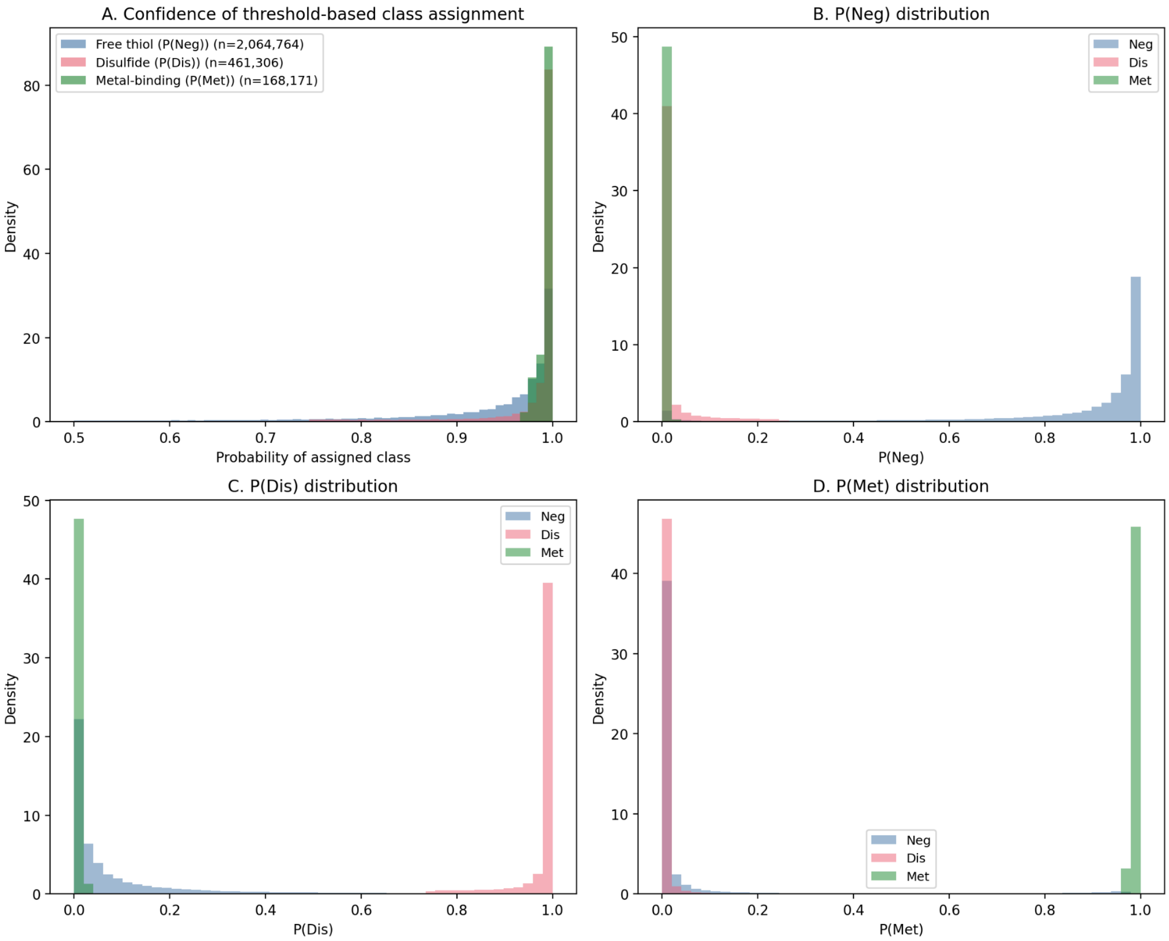


**Figure S3. (A)** Distribution of classification confidence (probability of the assigned class) for cysteines passing the operational thresholds. Disulfide assignments have median P(Dis) = 0.997; metal-binding assignments (n = 168,171) have median P(Met) = 0.998; free-thiol cysteines (n = 2,064,764) have median P(Neg) = 0.961. The thresholded positive classes are assigned with near-certainty, while the free-thiol class represents the absence of a specific functional signal and is correspondingly broader. **(B-D)** Per-class probability distributions colored by the threshold-assigned class, showing clear separation between classes with minimal overlap above the operational cutoffs.
